# A hierarchical model of perceptual multistability involving interocular grouping

**DOI:** 10.1101/800219

**Authors:** Yunjiao Wang, Zachary P Kilpatrick, Krešimir Josić

## Abstract

Ambiguous visual images can generate dynamic and stochastic switches in perceptual interpretation known as perceptual rivalry. Such dynamics have primarily been studied in the context of rivalry between two percepts, but there is growing interest in the neural mechanisms that drive rivalry between more than two percepts. In recent experiments, we showed that split images presented to each eye lead to subjects perceiving four stochastically alternating percepts (Jacot-Guillarmod et al., 2017): two single eye images and two interocularly grouped images. Here we propose a hierarchical neural network model that exhibits dynamics consistent with our experimental observations. The model consists of two levels, with the first representing monocular activity, and the second representing activity in higher visual areas. The model produces stochastically switching solutions, whose dependence on task parameters is consistent with four generalized Levelt Propositions. Our neuromechanistic model also allowed us to probe the roles of interactions between populations at the network levels. Stochastic switching at the lower level representing alternations between single eye percepts dominated, consistent with experiments.

## 1. Introduction

When conflicting images are presented to different eyes, our visual system often fails to produce a stable fused percept. Instead, perception stochastically alternates between the presented images (Wheatstone, 1838; Levelt, 1965; Leopold and Logothetis, 1999; Blake and Logothetis, 2002; Blake, 2001). More generally, multistable binocular rivalry between more than two percepts can occur when images presented to each eye can be partitioned and regrouped into coherent percepts. For example, subjects presented with the jumbled images in Fig. 1A may alternatively perceive a monkey face, or the jungle scene shown in Fig. 1B (Kovacs et al., 1996). In these cases perception evolves dynamically under constant stimuli, revealing aspects of the cortical mechanisms underlying visual awareness (Leopold and Logothetis, 1999; Tong et al., 2006; Sterzer et al., 2009; Leopold and Logothetis, 1996; Polonsky et al., 2000).

**Figure 1:**
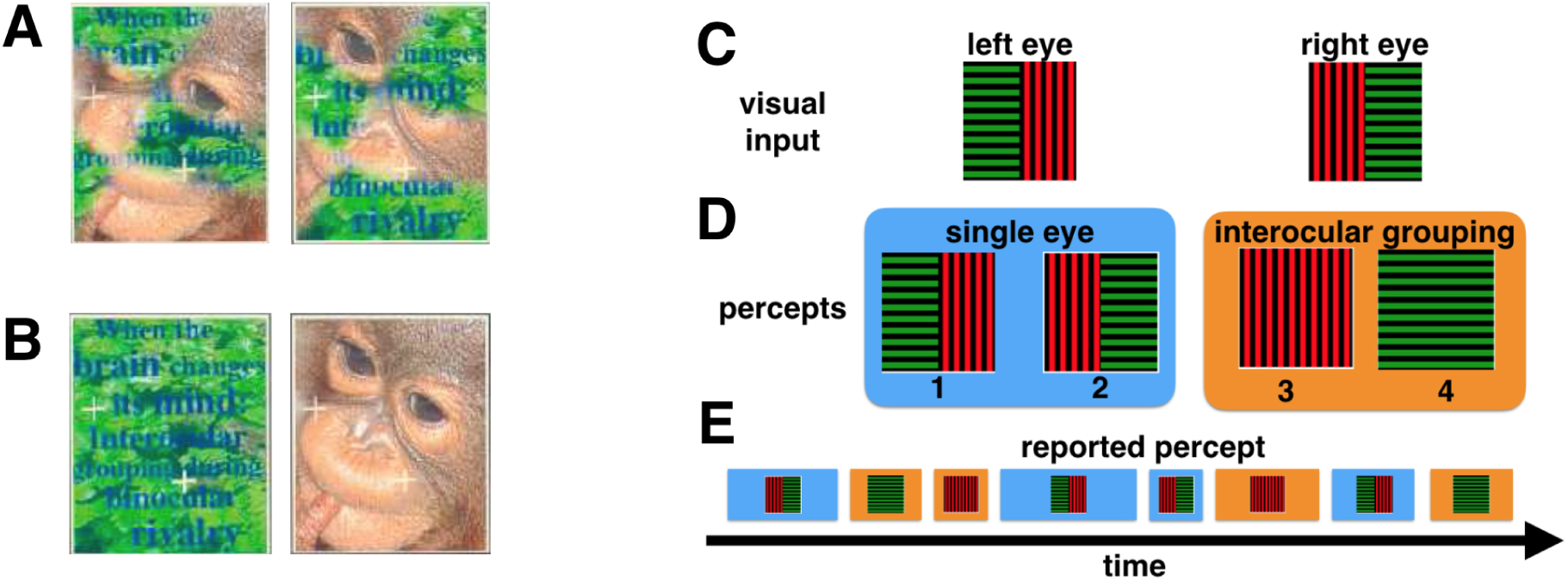
Multistable perceptual rivalry. The fragmented images presented to the left and right eyes in (A) can lead to the coherent percepts shown in (B) (Kovacs et al., 1996). (C) An example of the stimuli presented to the left and right eyes in Jacot-Guillarmod et al. (2017). Gratings were always split so that halves with the same color and orientation could be matched via interocular grouping, but were otherwise randomized across trials and blocks (See Jacot-Guillarmod et al. (2017) for experimental methods). (D) Subjects typically reported seeing one of four percepts – two single-eye and two grouped – at any given time during a trial. (E) A typical perceptual time series reported by a subject, showing the stochasticity in both the dominance times and the order of transitions between percepts.

While the literature on bistable binocular rivalry is extensive, far fewer studies have addressed multistable percepts. Rivalry between multiple percepts likely involves higher level image recognition, as well as monocular competition (Kovacs et al., 1996; Suzuki and Grabowecky, 2002; Huguet et al., 2014; Golubitsky et al., 2019), suggesting a noninvasive way to probe perceptual mechanisms across cortical areas, and offering a broader picture of visual processing.

Here, we build on previous models to provide a mechanistic account of perceptual multistability due to interocular grouping effects (Laing and Chow, 2002; Wilson, 2003; Moreno-Bote et al., 2007; Shpiro et al., 2007; Said and Heeger, 2013; Dayan, 1998). We propose a mechanism that involves different levels of visual cortical processing by building a hierarchical neural network model of binocular rivalry with interocular grouping. Our model captures the dynamics of perceptual switches reported by human subjects in experiments described by Jacot-Guillarmod et al. (2017) involving the visual stimuli shown in Fig. 1C. When presented with these stimuli, subjects reported alternations between four percepts, two *single-eye percepts*, and two *grouped percepts* that combine two halves of each stimulus into a coherent whole (See Fig. 1D).

Levelt’s four propositions (Levelt, 1965) capture the hallmarks of bistable binocular rivalry by relating *stimulus strength* (such as contrast or luminance), *dominance duration* (the time interval during which a single percept is reported), and *predominance* (the fraction of the time a percept is reported). Jacot-Guillarmod et al. (2017) have provided experimental support for a generalized version of Levelt’s propositions, and our model suggests neural mechanisms that drive the underlying cortical dynamics encoding perceptual changes.

Levelt’s propositions describe well–tested statistical properties of perceptual alternations (Laing and Chow, 2002; Brascamp et al., 2006; Wilson, 2007; Moreno-Bote et al., 2010; Klink et al., 2010; Seely and Chow, 2011), and provide constraints on mechanistic models of binocular rivalry. Successful models broadly explain rivalry in terms of three interacting neural mechanisms: *Mutual inhibition* drives the exclusivity of the perceived patterns; *Slow adaptation* drives the transition between the different percepts; Finally, *internally generated noise* is necessary to account for the observed variability in perceptual switching times (Matsuoka, 1984; Lehky, 1988; Arrington, 1993; Lumer, 1998; Kalarickal and Marshall, 2000; Laing and Chow, 2002; Lago-Fernandez and Deco, 2002; Stollenwerk and Bode, 2003; Wilson, 2003; Noest et al., 2007; Seely and Chow, 2011; Freeman, 2005; Brascamp et al., 2006; Moreno-Bote et al., 2007).

In our model we includes these mechanisms, along with additional, abstracted features of the visual system. The model contains a lower level associated with early (e.g., eye-based) neural processes and tuned to geometric stimulus properties (e.g. orientation), and a higher level which accounts for complex pattern grouping and is responsible for the formation of late stage percepts. Our model thus extends earlier models of bistable binocular rivalry, and it reduces to simpler rivalry models under bistable inputs (Wilson, 2003; Tong et al., 2006; Diekman et al., 2013).

We hypothesize that pattern grouping effects occur already at the early stages of the visual system. We thus assume that the connectivity of the earlier, first layer in our network is modulated by cues – in our case color saturation – indicating which parts of the percepts belong to the same group. In most previous models of rivalry the strength of the stimulus primarily modulated the inputs to various network modules. In our case, we assume that input strength changes the connectivity between the neural populations at the lower level of the network.

We found that our model captured the statistics of perceptual alternations reported by experimental subjects Jacot-Guillarmod et al. (2017). Moreover, over a range of parameters the model also displays dynamics consistent with the generalized version of Levelt’s Propositions proposed by Jacot-Guillarmod et al. (2017). Our results hold under weak feedback from the higher level to the lower level. However, we observed these dynamics only with strong mutual inhibition between populations representing conflicting stimuli at the lower level of the visual hierarchy. Our model thus suggests constraints on the interactions between neural populations in the visual system consistent with experimentally observed perceptual dynamics.

Our study thus shows that more complex visual stimuli can be used in perceptual rivalry experiments to drive the development of more detailed mechanistic models of perceptual processing (Wilson, 2003; Dayan, 1998; Freeman, 2005).

## 2. Methods

### 2.1. Hierarchical model of perceptual multistability with interocular grouping

Considerable evidence suggests that visual processing in humans and other mammals is organized hierarchically (Polonsky et al., 2000; Tong, 2001; Leopold and Logothetis, 1996; Logothetis and Schall, 1989; Sheinberg and Logothetis, 1997; Dayan, 1998; Wilson, 2003; Freeman, 2005; Tong et al., 2006). The simplest models of such processing assume that visual areas at the higher level of the hierarchy pool the activity of lower areas (Riesenhuber and Poggio, 1999). Here we extend previous, non-hierarchical models of perceptual rivalry (Laing and Chow, 2002; Wilson, 2009; Moreno-Bote et al., 2007; Huguet et al., 2014; Diekman et al., 2013) to a model that spans two levels of the visual hierarchy, and study grouping in perceptual competition. A schematic representation of our model is shown in Fig. 2. The sub-network at the first level of the hierarchy consists of four neural populations, each receiving input from a different hemifield of the two eyes (See also Fig. 6C of Diekman et al. (2013) and Fig. 2B of Tong et al. (2006)). The responses of all four possible pairs of populations at the first level are integrated by distinct populations at the second level (Laing and Chow, 2002; Wilson, 2003; Moreno-Bote et al., 2007). Each of the four populations at the second level corresponds to one of the four percepts shown in Fig. 1B.

**Figure 2:**
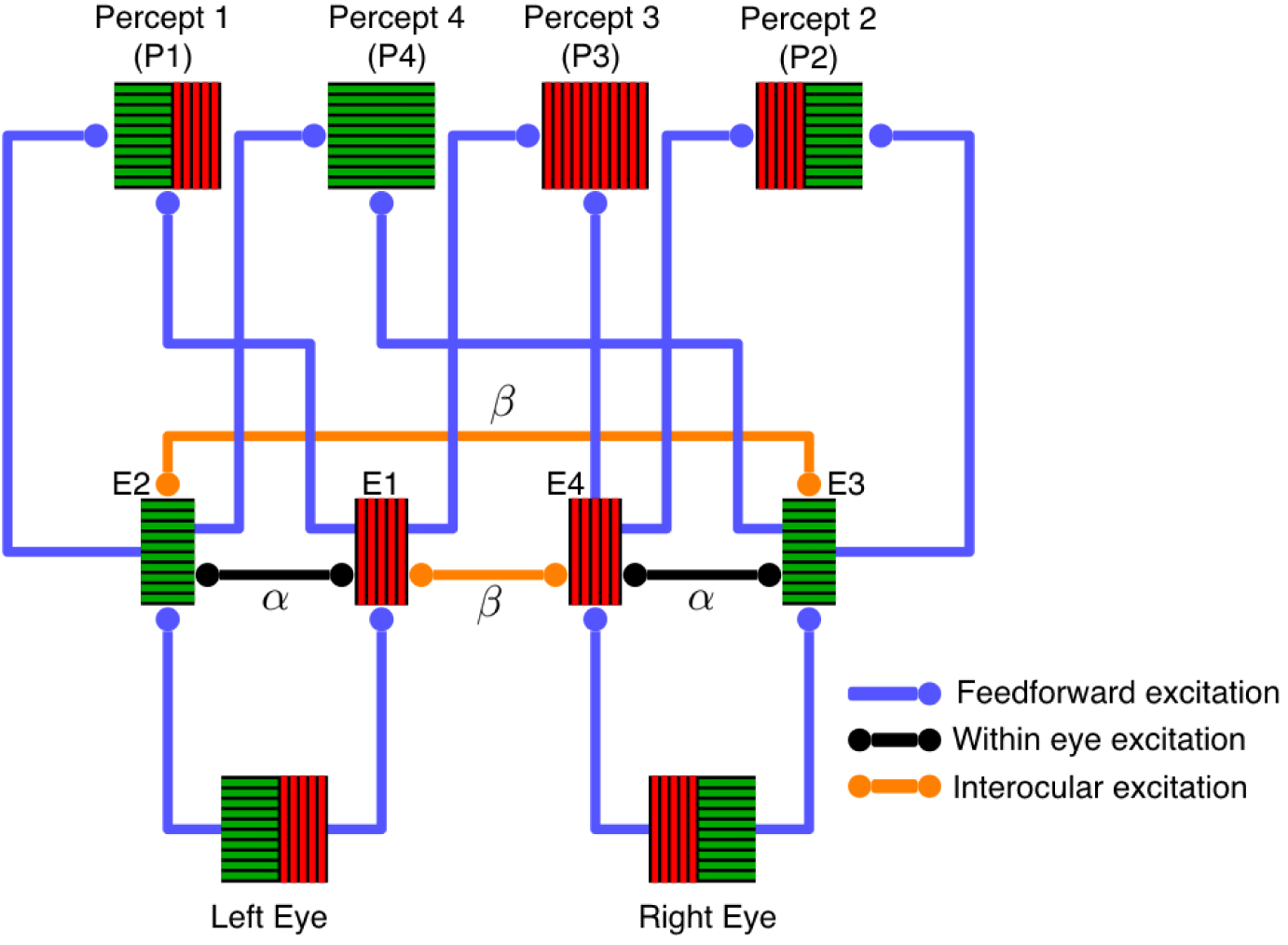
A hierarchical model of interocular grouping. Neural populations representing stimuli to the four hemifield-eye combinations at Level 1 provide feedforward input to populations representing integrated percepts at Level 2, as described by Eqs. (1) and (4) (See also Fig. 6C of Diekman et al. (2013) and Fig. 2B of Tong et al. (2006)). The figure shows recurrent excitation within Level 1. To avoid clutter, mutual inhibition between the same hemifield of opposite eyes is not shown. All populations at the second level of the hierarchy mutually inhibit one another (Laing and Chow, 2002; Wilson, 2003; Moreno-Bote et al., 2007).

**Figure 3:**
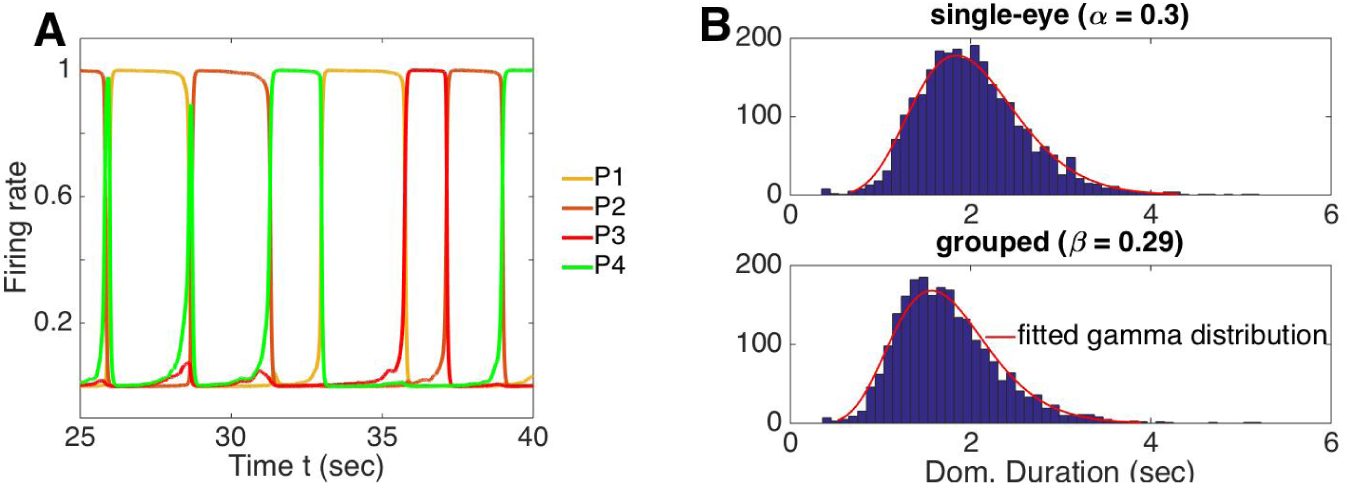
Dynamics of a hierarchical model of interocular grouping. (A) A typical time series of the firing rates, *P*_*i*_, of neural populations at the second level of the model. Each of these populations is associated with one of the four percepts: *P*_1_ and *P*_2_ correspond to single-eye percepts, and *P*_3_ and *P*_4_ correspond to grouped percepts. Here we used same-eye coupling *α* = 0.3, interocular grouping strength *β* = 0.26, and input strength *I*_*i*_ = 1. (B) Distributions of dominance durations in the model have a single mode around 1.8s for single-eye percepts, and 1.5s for grouped percepts. These distributions are consistent with experimental data. Distributions were obtained from 100 time series, each 100*s* in duration. Parameters were set to *I*_*i*_ = 1.2, *w* = 1, *g* = 0.5, *c*_*i*_ = 1, *v* = *γ* = 0.45, *κ* = 0.5.

**Figure 4:**
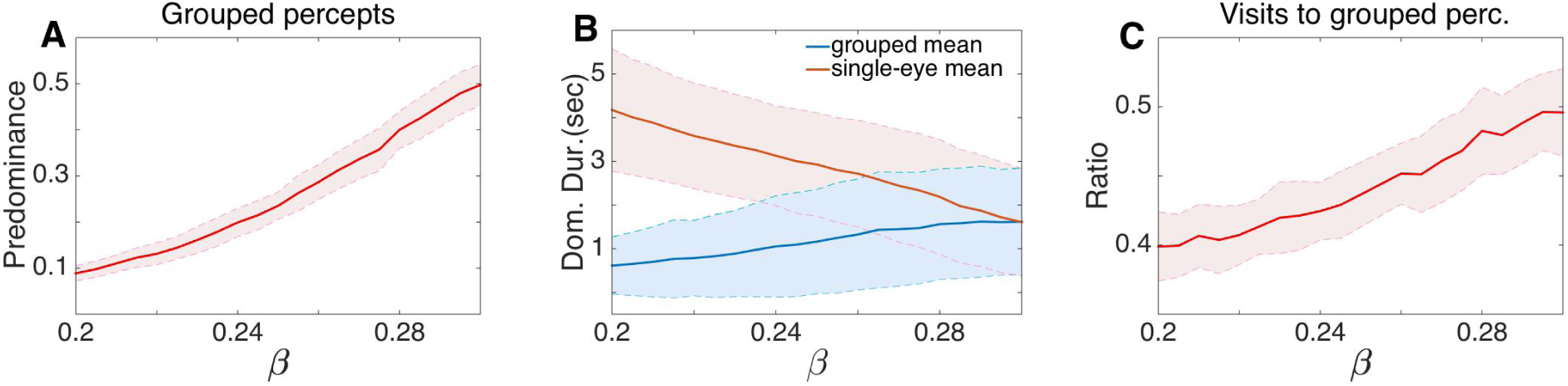
Effects of varying the interocular grouping strength, *β*, at the first level of the hierarchical model. (A) Predominance of grouped percepts increased with *β*. (B) The average dominance duration of single-eye percepts decreased with *β*, while that of grouped percepts remained approximately unchanged, particularly in the range 0.27 *≤ β ≤* 0.3. (C) Furthermore, the frequency of visits to grouped percepts increased with *β*. Other parameters were the same as in Fig. 3. Solid lines represent computationally obtained means, and shaded regions represent one standard deviation about the means obtained over 100 realizations.

**Figure 5:**
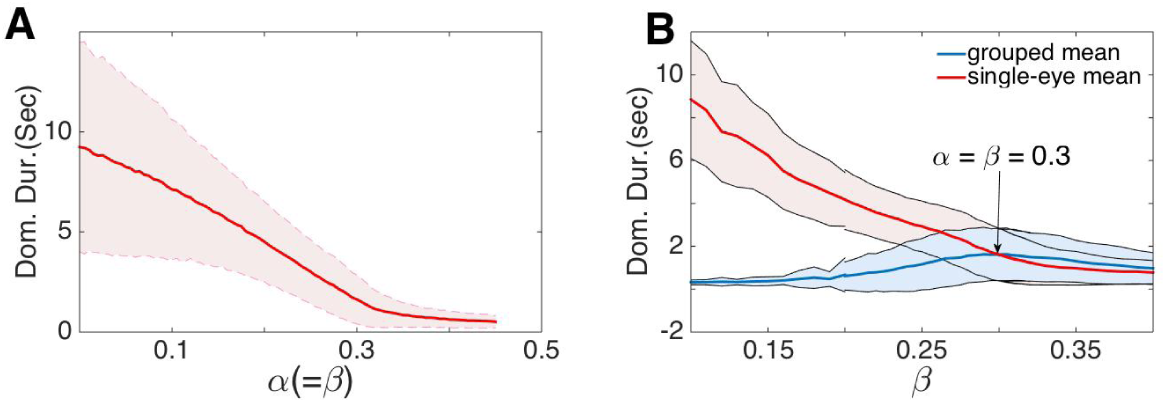
Levelt’s Proposition IV holds in the hierarchical model. (A) Increasing within- and between-eye grouping strengths (*α* and *β* respectively), simultaneously while keeping them equal decreased the average dominance duration. (B) Proposition II held when *β < α*. Here *α* = 0.3, with other parameter values as in Fig. 3.

**Figure 6:**
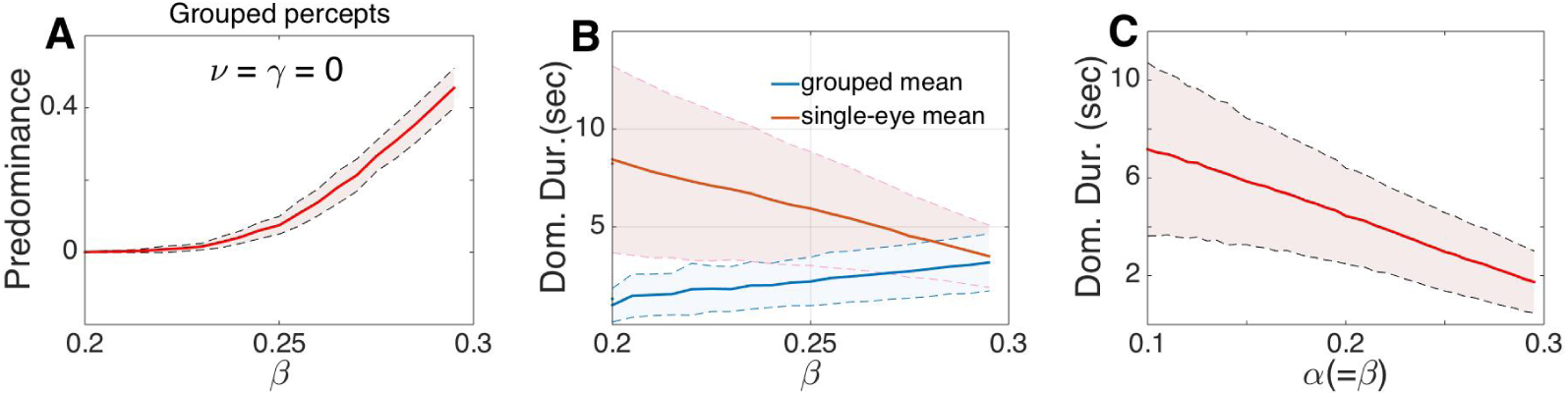
Levelt’s propositions hold without mutual inhibition at Level 2 (*v* = *γ* = 0). (A) Predominance of grouped percepts increased with interocular grouping strength, *β*. (B) The average dominance duration of single-eye percepts (stronger percepts) decreased much faster than the average dominance duration of grouped percepts (weak percepts). (C) The average dominance duration decreased as *α* and *β* were increased and kept equal. Other parameter values as in Fig. 3.

A key feature of our model is the presence of *excitatory* coupling between populations receiving input from different hemifields both from the same and from different eyes. This is consistent with electrophysiology and tracing experiments that reveal long-range horizontal connections between neurons in area V1 with non-overlapping receptive fields, but similar orientation preferences (Stettler et al., 2002; Sincich and Horton, 2005). We also assumed *inhibitory* coupling between populations receiving conflicting input from the same hemifield of different eyes, *e.g*. the left hemifield of the left and the left hemifield of the right eye. Experimental literature suggests cells with orthogonal orientation preferences can inhibit one another through multisynaptic pathways involving recurrent and feedback circuitry (Ringach et al., 1997; Ferster and Miller, 2000). Finally, we assumed that all populations at the second level inhibit each other, as in previous computational models (Laing and Chow, 2002; Moreno-Bote et al., 2007; Shpiro et al., 2007; Lankheet, 2006; Seely and Chow, 2011).

The two levels thus form a processing hierarchy (Wilson, 2003; Tong et al., 2006) with the first roughly associated with monocular neural activity generated in LGN and V1 (Wilson, 2003; Blake, 1989; Polonsky et al., 2000; Tong, 2001), and the second level associated with the activity of higher visual areas, such as V4 and MT, that process objects and patterns (Leopold and Logothetis, 1999; Wilson, 2003; Lamme and Roelfsema, 2000). However, each level could also describe multiple functional layers of the visual system (Sterzer et al., 2009).

#### First level of the visual hierarchy

The activity of each neural population receiving input from one of the four hemifield-eye combinations at Level 1 is described by a firing rate variable *E*_*i*_, *i* = 1, 2, 3, 4 (corresponding to left hemi/left eye; right hemi/left eye; left hemi/right eye; and right hemi/right eye, see Fig. 2). To model adaptation, we included variables describing hyperpolarizing currents activated at elevated firing rates, *H*_*i*_, with *i* = 1, 2, 3, 4 (Benda and Herz, 2003). The firing rates at the lower level of the visual hierarchy are then governed by the following equations:

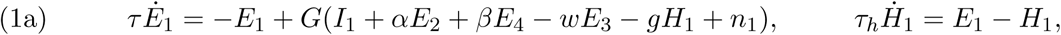

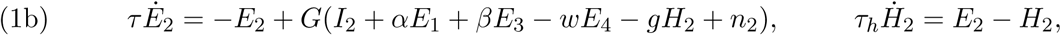

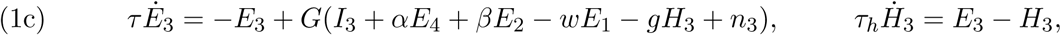

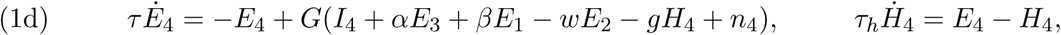

with activity time constant *τ* = 10ms (Häusser and Roth, 1997) and adaptation time constant *τ*_*h*_ = 1000ms. The inputs, *I*_*i*_, model the strength of the stimulus in each hemifield, and *g* is the strength of adaptation. We assumed that all inputs, *I*_*i*_, all are equal in intensity, so that *I*_*i*_ = *I* for *i* = 1, 2, 3, 4. This is consistent with the experiments of Jacot-Guillarmod et al. (2017) where stimuli were calibrated to be equal in intensity.

The strength of within eye excitatory coupling is determined by the parameter *α*, while interocular excitatory coupling between populations receiving input from complementary hemifields is described by *β*. The strength of mutual inhibition due to orientation and color competition is determined by *w*.

We used a sigmoidal gain function, *G*(*x*), to relate the total input to the population to the output firing rate,

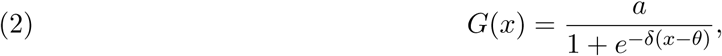

where *a* = 1, *δ* = 10 and *θ* = 0.2. This choice was not essential, as we could have used other gain nonlinearities, such as a Heaviside step or a rectified square root, as long as each individual population, *E*_*i*_, has a bistable regime (with a low and high stable firing rate state) for a given input *I*_*i*_ (Laing and Chow, 2002; Moreno-Bote et al., 2007).

Random fluctuations due to network effects and synaptic noise were modeled by the variables *n*_*i*_, *i* = 1, 2, 3, 4 (Faisal et al., 2008). Following Moreno-Bote et al. (2007), we modeled the fluctuations in the total input to each population as an Ornstein-Uhlenbeck process,

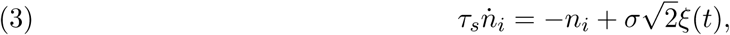

where *τ*_*s*_ = 200*ms, σ* = 0.03, and *ξ*(*t*) is a white-noise process with zero mean. Changing the timescale and amplitude of noise does not impact the results significantly.

#### Second level of the visual hierarchy

As shown in Fig. 2, feedforward connectivity from Level 1 to Level 2 of the network associates each of four possible combinations of hemifields with a distinct percept reported by obsevers, and a distinct population at the second level of the hierarchy. The activity of each of these populations is governed by the firing rate, *P*_*i*_, and an associated adaptation variable, *A*_*i*_, *i* = 1, 2, 3, 4,

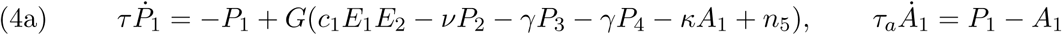

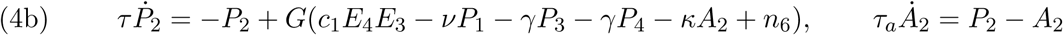

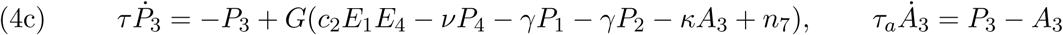

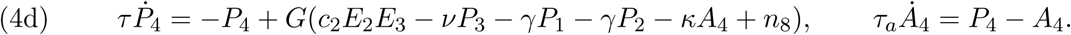

For simplicity we assumed that the activation rate, *τ*, and adaptation rate, *τ*_*a*_ *≡ τ*_*h*_ are equal between layers.

Feedforward inputs to the second level were modeled as a product of activities of the associated populations at the first level. For instance, population activity *P*_1_ depends on the product *E*_1_*E*_2_ since Percept 1 is composed of the two stimuli in the same-eye hemifields providing input to populations 1 and 2 at Level 1 (*e.g*. the horizontal green bar, and vertical red bar presented to the left eye in the example shown in Fig. 2). Experimental and modeling studies have pointed to such multiplicative combinations of visual field segments as a potential mechanism for shape selectivity (Salinas and Abbott, 1996; Brincat and Connor, 2006). When we replaced the multiplicative input to the second level population with additive input from Level 1, *E*_*j*_ + *E*_*k*_, our results remained qualitatively similar.

#### Feedback from upper-level

Experimental results suggest that top-down processing can influence rivalry (Bartels and Logothetis, 2010; Klink et al., 2008). We have thus also considered an extension of our model by that includes feedback from Level 2 to Level 1,

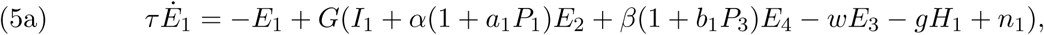

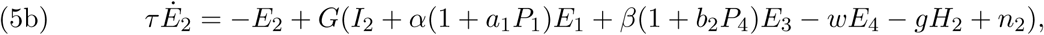

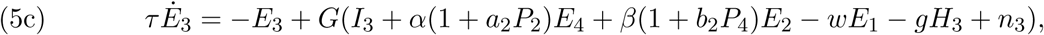

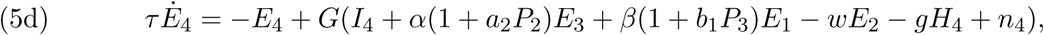

We compare the dynamics of the networks with and without feedback, and discuss the impact of feedback in Results.

### 2.2. Parameter Values

As with many previous models of rivalry, the dynamics of our model depends on the choice of parameters, but is relatively robust: We set the time scales, *τ*, *τ*_*h*_ and *τ*_*s*_, to values found in computational modeling studies and suggested by experimental work neural population activity dynamics, spike frequency adaptation, and temporal correlations in population-wide fluctuations (Häusser and Roth, 1997; Benda and Herz, 2003; Moreno-Bote et al., 2007; Renart et al., 2010). Other parameters were first chosen so that in the absence of noise the model displayed periodic solutions corresponding to alternations of single-eye percepts. We then included noise, and searched for parameters that produced dynamics that agreed with experimental results. For more details, see Appendix A and Fig. 10 therein.

**Figure 7:**
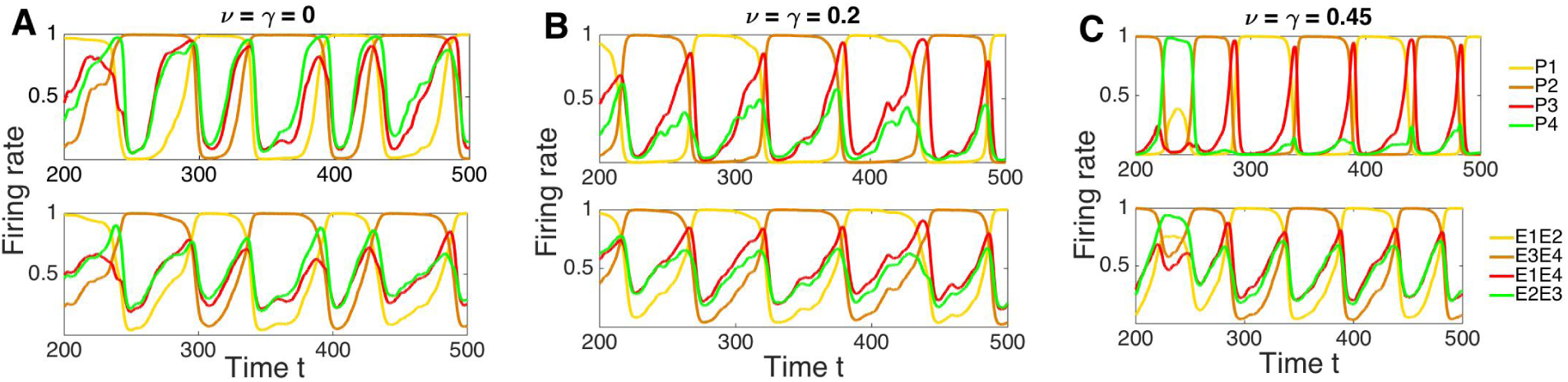
Time series with different mutual inhibition at the upper level. Each upper panel shows the neural activity of percepts (populations at the higher level of the hierarchy), and lower panels show inputs from the lower to the higher level of the hierarchical model; e.g., *E*1*E*2 is the input to *P*1. (A,B) Weak or mild mutual inhibition at the higher level helped disentangle different percepts, i.e. mutual inhibition at the upper level increased the distance between the activity levels of the dominating percepts and suppressed percepts; whereas (C) strong inhibition at the higher level lead to more frequent percept switching. Other parameter values as in Fig. 3.

**Figure 8:**
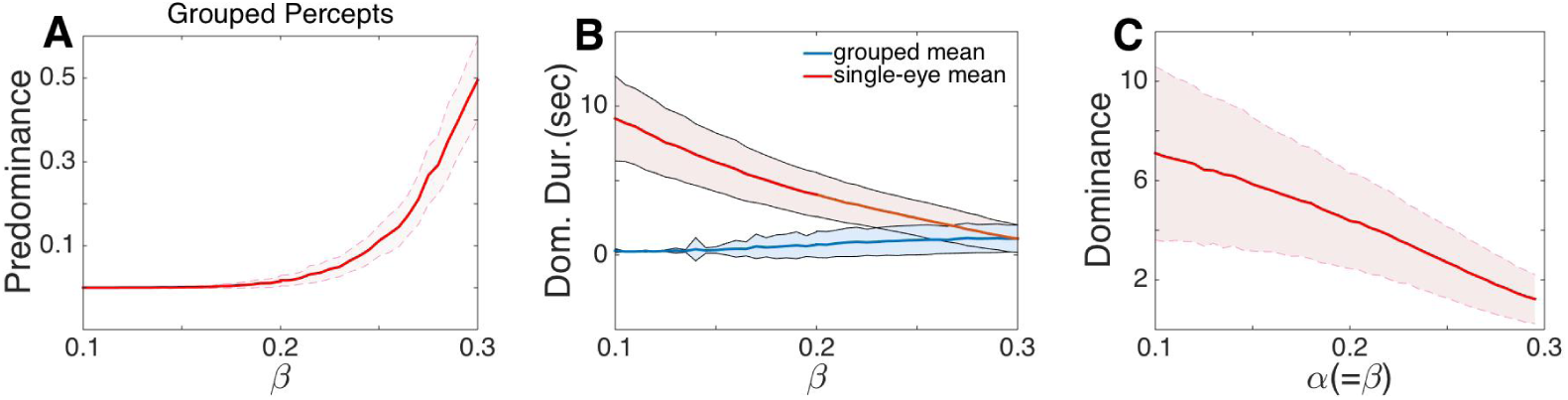
Generalized Levelt’s propositions hold in the absence of adaptation at the higher level of the visual hierarchy. (A) The predominance of grouped percepts increased with the interocular grouping strength, *β*. (B) The average dominance duration of single-eye percepts (stronger percepts) decreased much faster than the average dominance duration of grouped percepts (weak percepts). (C) The average dominance duration decreased with *α* and *β* when the two were kept equal. Parameter values as in Fig. 3.

**Figure 9:**
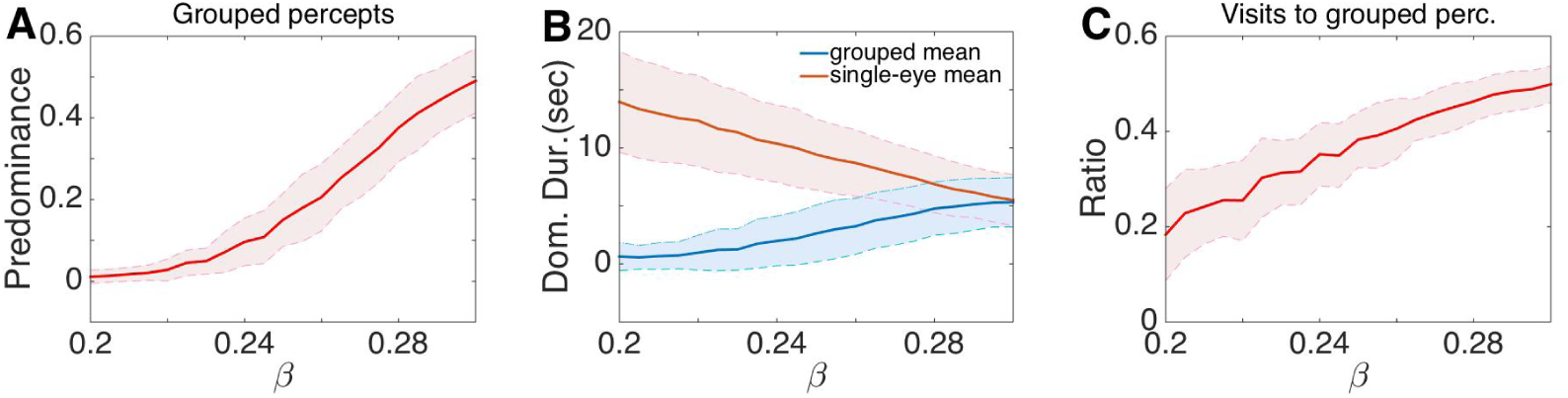
Simulation results with feedback from the higher to the lower level of the hierarchy. Simulations indicate that the model can capture the key experimental results in (Jacot-Guillarmod et al., 2017) even with feedback from the higher level to the lower level: (A) Pre-dominance of grouped percepts increased as the interocular grouping strength increased; (B) The average dominance duration of single-eye percepts decreased while the average dominance duration of grouped percepts remained approximately unchanged (when *β < α* but close to the value *α*); (C) The ratio of the number of visits to the grouped percepts increased as the interocular grouping strength increased. Here *a*_*i*_ = *b*_*i*_ = 0.1 in (5), with other parameters as in Fig. 3.

**Figure 10:**
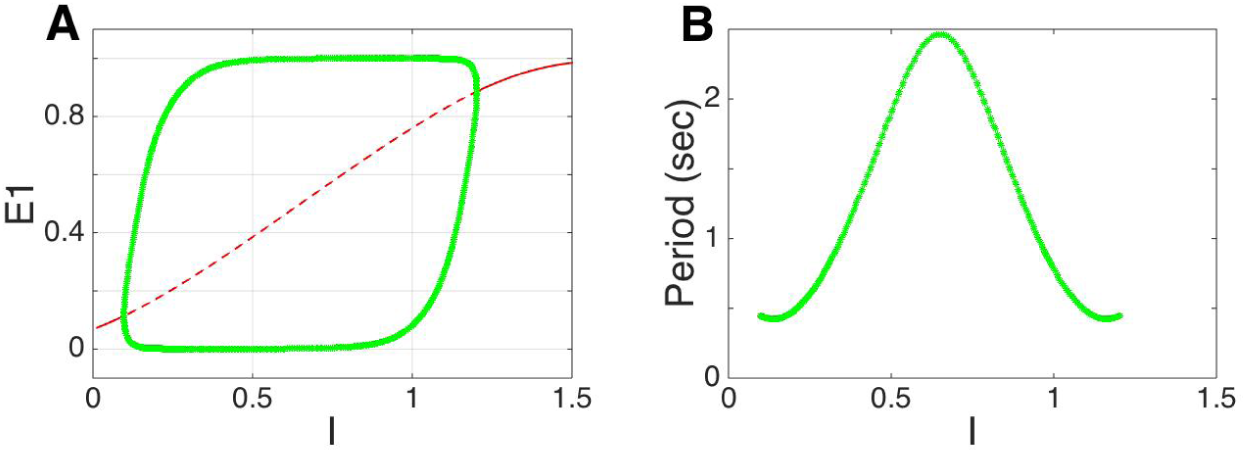
The hierarchical model captures conventional bistable binocular rivalry. (A) The bifurcation diagram with bifurcation parameter *I* when *α* = *β* = 0.3, and other parameters as in Fig. 3 shows the emergence and disappearance of periodic solutions. The green curves represent the branches of a stable periodic solution, the solid red curve represents stable equilibria, and the dashed red curve represents unstable equilibria; (B) The period of the corresponding stable periodic solution peaks around *I* = 0.6.

## 3. Results

We use the hierarchical model described by Eqs. (1) and (4) to explain the different experimentally observed features of binocular rivalry involving interocular grouping. Moreover, we show that our model can provide a unifying mathematical framework that accounts for the generalized Levelt’s propositions, and provides concrete hypotheses of how different neural mechanisms shape perceptual dominance across levels of the visual hierarchy. At the same time, our model reduces to previous successful models of binocular rivalry with stimuli that conflict between the eyes, but do not allow inter-ocular grouping. We use numerical experiments and bifurcation theory to demonstrate the qualitative changes in the dynamics of the model to support these conclusions.

### 3.1. Levelt’s Propositions and their generalization

Levelt’s propositions relate stimulus strength to *dominance duration* – the time interval during which a single percept is reported; *predominance* – the fraction of the time a percept is reported; and *alternation rate* – the rate of switching between percept reports. In the context of bistable rivalry, Levelt’s propositions have most recently been stated as (Brascamp et al., 2015): (I) Increasing the strength of the stimulus presented to one eye increases the predominance of that stimulus; (II) Increasing the difference in stimulus strengths between the two eyes increases the dominance duration of the stronger stimulus; (III) Increasing the difference in stimulus strengths between the two eyes reduces the perceptual alternation rate; (IV) Increasing stimulus strength in both eyes while keeping it equal between eyes increases the perceptual alternation rate. This effect may reverse at near-threshold stimulus strength (See Fig. 3 in Brascamp et al. (2015) for an illustration).

The *strength of a percept* has been defined as any attribute whose increase causes that percept to suppress the appearance of other percepts (Brascamp et al., 2015). Levelt’s Proposition I thus effectively defines the strength of a percept attribute according to whether it impacts a percept’s predominance. Jacot-Guillarmod et al. (2017) found experimental support for some of the following extensions of Levelt’s proposition using the stimuli and associated percepts shown in Fig. 1C,D:

I. *Increasing percept strength of grouped percepts or single-eye percepts increases the perceptual predominance of those percepts*. Jacot-Guillarmod et al. (2017) showed that increasing color saturation increases the predominance of grouped percepts. Experimental results thus support this proposition, with color saturation defining the strength of the grouped percept class.
II. *Decreasing the difference between the strength of the grouped percepts and that of single-eye percepts primarily decreases the average dominance duration of the stronger percepts*. When the single-eye percept is stronger (weaker), increasing the strength of grouped percepts decreases (increases) the average dominance duration of the single-eye (grouped) percepts. Jacot-Guillarmod et al. (2017) showed that increasing color saturation primarily decreased the average dominance duration of the stronger, single-eye percepts, consistent with Proposition II. Experimental results did not speak to the validity of generalized Proposition II when the grouped percepts were stronger. When one class of percept is much stronger (e.g., single-eye percepts), we expect them to completely suppress percepts of the other class (e.g., grouped percepts). Percept strengths used in the experiments of Jacot-Guillarmod et al. (2017) were not sufficiently high to validate these predictions, but we test them in our model.
III. *Decreasing the difference in strengths between grouped percepts and single-eye percepts increases the perceptual alternation rate*. Since alternation rate and average dominance duration are related reciprocally, Proposition III follows from Proposition II.
IV. *Increasing the strength in both grouped percepts and single-eye percepts while keeping strength equal among percepts increases the perceptual alternation rate*. Proposition IV was not tested directly in Jacot-Guillarmod et al. (2017), as changing color saturation affected the strengths of each percept differently. We show below that this Proposition holds in our model.

### 3.2. The hierarchical model exhibits perceptual multistabiliy

We first demonstrate how our model captures alternations between multiple percepts. As in previous studies, we associated a neural population with each percept: An elevation in the activity of a population at Level 2 of our model indicates that the corresponding percept is perceived and reported (Laing and Chow, 2002; Wilson, 2003; Moreno-Bote et al., 2007; Dayan, 1998; Freeman, 2005; Wilson, 2009; Lehky, 1988; Said and Heeger, 2013; Lago-Fernandez and Deco, 2002; Lumer, 1998).

For a wide range of parameters, a single Level 2 neural population exhibited elevated activity, and suppressed the activity of the remaining populations (See Fig. 3A for a representative simulation). The order and timing of these periods of elevated firing were stochastic, and the distributions of the time periods of elevated firings were unimodal (Fig. 3B). This dynamics corresponded to the reports of experimental subjects who primarily reported seeing individual percepts over intervals of varying durations, and random alternations between the percepts. Consistent with previous models (Laing and Chow, 2002; Wilson, 2003; Moreno-Bote et al., 2007), stochastic alternations between percepts emerged due to the mutual suppression between the four populations at the second level of the hierarchy, while noise and adaptation drove alternations between the active populations.

### 3.3. Changing stimulus strength in the model yields experimentally observed dominance duration changes

In classical models of perceptual rivalry, stimulus and percept strengths are represented by the magnitude of input(s) to different neural populations. Changes in these input strengths correspond to changes in stimulus features like luminosity or contrast (Freeman, 2005; Seely and Chow, 2011). In the case of rivalry with grouped percepts (Fig. 1D), we assume that changes in color saturation have little effect on the strength of the inputs *I*_*i*_ (Jacot-Guillarmod et al., 2017). Rather, we assume that varying color saturation changes the tendency for interocular grouping between the two halves of images of the same color and orientation, consistent with Gestalt principles of similarity (Roelfsema, 2006; Kohler, 2015). Thus color saturation provides a visual cue for binding complementary halves of grouped percepts (Wagemans et al., 2012). We therefore modeled the effects of color saturation as a change in the strength of cross-hemispheric excitatory connections, *β*, between populations responding to like stimulus features. We also assumed that the excitatory coupling, *α*, between populations reprsenting same-eye image halves was unaffected by changes in color saturation.

Jacot-Guillarmod et al. (2017) made several observations about the impact of color saturation on perceptual alternations recapitulated by our model. First, color saturation increased subjects’ *predominance* of grouped percepts, *i.e*. the fraction of the total time subjects reported a grouped percept out of the total time they reported seeing any percept: Increasing interocular coupling strength, *β*, in our model also increased the predominance of grouped percepts (See Fig. 4A). Thus color saturation, modeled by connection strength, *β*, between first level network populations in our model, satisfies the commonly used definition of *stimulus strength* (Brascamp et al., 2015).

Second, Jacot-Guillarmod et al. (2017) observed that increasing color saturation decreased the average dominance duration (the average time the percept is seen before a switch occurs) of single-eye percepts while the average dominance duration of grouped percepts remained largely unchanged. Our model captured this feature over a range of parameters: For 0.2 *< β <* 0.3, increasing *β* decreased the dominance duration of single-eye percepts, while changes in dominance of grouped percepts were smaller and nearly absent as *β* approached *α* (See Fig 4B).

Finally, Jacot-Guillarmod et al. (2017) showed that increasing color saturation increased the ratio of visits to grouped percepts. Our model exhibits this behavior as well: The ratio of visits to grouped percepts increased with interocular grouping strength, *β*, (See Fig 4C). As shown in Fig. 9 these results also hold in the presence of feedback.

### 3.4. Our model conforms to the generalized Levelt’s propositions when *α > β*

We next asked whether the dynamics of our model agrees with experimentally observed generalizations of Levelt’s propositions (Jacot-Guillarmod et al., 2017). As shown in Fig. 4, Proposition I hold. In fact, the proposition holds over a wide range of parameter values, even when other propositions fail, and in all model versions we have explored.

We found that Proposition II holds in our model when *β < α*. When excitatory coupling between neural populations representing different-eye hemispheres is weaker than coupling between same-eye hemisphere populations, increasing interocular coupling strength *β* decreases the average dominance duration of the two single-eye percepts but very weakly increases the average dominance duration of the grouped percepts (See Fig. 5B). Since Proposition III follows from Proposition II and I, our model supports Proposition III as well.

To determine whether our model conforms to the prediction of Proposition IV, we varied *α* and *β* simultaneously while keeping them equal (See Fig. 5A). When grouping strength, *β*, is sufficiently high (*β >* 0.32), multiple subpopulations become co-active, indicating fusion. Fig. 5A shows that an increase in *β* (and *α*) decreased the average dominance duration of both grouped and single-eye percepts, *i.e*. increasing the strengths of all percepts while keeping them equal increases the perceptual switching rate, in accord with Proposition IV. As in existing models for bistable binocular rivalry, Levelt’s Propositions IV holds only for parameter values over which the period of the periodic solutions of the associated deterministic model decreases as *I* increases (See Fig. 10 in Appendix for more details).

#### Remark

To explore the full range of model behaviors, we also consider the case *α < β* representing strong interocular coupling. In this case, Proposition II fails since increasing the strength of the grouped percepts by increasing *β* does not lead to an increase in their average dominance duration, despite the grouped percepts being stronger (Fig. 5B). Such failures are common in other existing models when percept strengths are close (See Fig. 11C which reproduces results from Seely and Chow (2011)). Proposition II states that the average dominance duration of the stronger percept should change more than that of the weaker percept, but this effect does not hold when input strengths are close in mutual inhibitory models of perceptual bistability (Fig. 11C and (Seely and Chow, 2011)).

**Figure 11:**
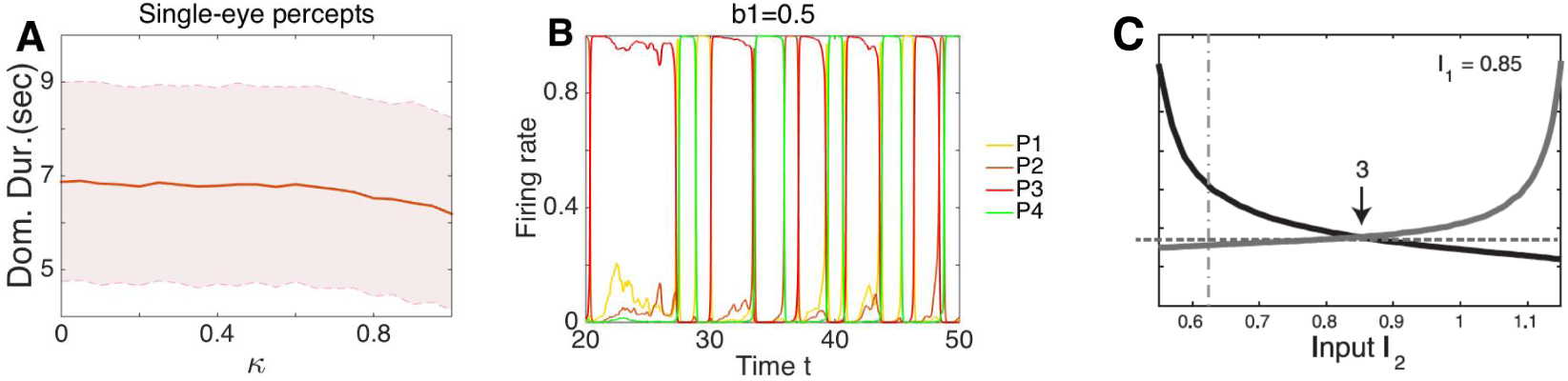
Adaptation rate, *κ*, at the higher level of the hieararchy, and top-down influence. (A) The adaptation rate had little or no effect on the dominance duration of percepts. Parameter values as in Fig. 3. (B) Example of top-down influence from only one percept, here *P*_3_ (*a*_1_ = *a*_2_ = *b*_2_ = 0 and *b*_1_ = 0.5). Top down input from one percept increased its dominance duration. Parameters not listed were as in Fig. 3. (C) Part of Fig. 4C from (Seely and Chow, 2011): Proposition IV did not hold when *I*_2_ ∈ (0.85, 1) since the increasing rate of the stronger percepts did not exceed the decreasing rate of the weak percept.

When the percept strength of the grouped percepts is much stronger than that of the single-eye percepts, perception is dominated by two rivaling grouped percepts. According to the original Levelt’s Proposition IV, further increasing in the strength of the grouped percepts should increase the switching rate between the two grouped percepts, reducing their average dominance duration. This is the case in our model, and is the reason for the decrease in average dominance duration when *β > α* shown in Fig. 5B, in contrast to the increase seen in Fig 11C).

### 3.5. The mechanisms of multistable rivalry in the hierarchical model

We next describe the mechanisms that drive the perceptual switching dynamics in our model. The neural interactions implied by these mechanisms may underlie the dynamics described by the generalized Levelt’s Propositions:

1. Increasing interocular grouping strength, *β*, promotes co-activation of populations *E*_1_ and *E*_4_, as well as *E*_2_ and *E*_3_ at the first level of the hierarchy. Joint activity of populations *E*_1_ and *E*_4_ leads to increased activation of population *P*_3_ at the second level. Similarly, joint activity of *E*_2_ and *E*_3_ increases activation of *P*_4_. Due to mutual inhibition between populations at the same hemifileds of opposite eyes, *E*_1_ and *E*_3_ (*E*_2_ and *E*_4_) synchronous activity of the pair *E*_1_ and *E*_4_ (*E*_2_ and *E*_3_) is likely not to be observed together with a coactivation of *E*_1_ and *E*_2_, or *E*_3_ and *E*_4_. Thus, a coactivation of the input *E*_1_*E*_4_ to *P*_3_ (*E*_2_*E*_3_ to *P*_4_) decreases the likelihood of elevated inputs *E*_1_*E*_2_ and *E*_3_*E*_4_ to the populations *P*_1_ and *P*_2_ corresponding to single-eye percepts. This explains why increasing interocular grouping strength, *β*, increases the predominance of the grouped percepts (*P*_3_ and *P*_4_), and hence the mechanism behind Proposition I.
2. As in earlier models of bistable rivalry, our hierarchical model exhibits perceptual switches either due to (a) inhibition release, or (b) escape driven by noise or the relaxation of adaptation (Curtu et al., 2008; Moreno-Bote et al., 2007). These two mechanisms are not mutually exclusive, and depend on model parameters. We chose parameters such that the escape mechanism dominates.
3. Keeping *α* = *β* and increasing their values is ‘equivalent’ to increasing the input, *I*: When single-eye percepts dominate, the two terms *αE*_2_ + *βE*_4_≈*α* in the gain of *E*_1_ in Eq. (1a). A similar observation applies to the corresponding two terms determining the evolution of the firing rates *E*_2_, *E*_3_ and *E*_4_, and a similar effect occurs when the grouped percepts dominate. Hence, simultaneously increasing the value of *α* and *β* while keeping them equal, is approximately equivalent to increasing the input *I*. Because of the choice of our parameter region in which the period of the associated deterministic model decreases as *I* increases, this implies that Proposition IV holds.

### 3.6. Impact of mutual inhibition at different levels of the hierarchical model

It has been debated at which level of the visual hierarchy mutually inhibitory interactions lead to rivalry (Carlson and He, 2004; Andrew and Lotto, 2004; Wilson, 2003). Carlson and He (2004) showed that incompatibilities (conflicting interocular information that cannot be fused) at the lower level are necessary for producing rivalry. In contrast, Andrew and Lotto (2004) used identical stimuli within a different chromatic surround to show that the presence of rivalry can depend on the perceptual meaning of the visual stimuli, and must thus occur at higher levels of the visual processing hierarchy. Wilson (2003), on the other hand, used a two-stage feedforward model to show that the elimination of mutual inhibition at early stages reveals the activity at the higher layer, *i.e*. the activity remains at steady-state at the first level, and rivalry occurs only at the higher level.

Our model exhibits behavior similar to that reported by Wilson (2003): If lower-level mutual inhibition is not strong enough, activity at the lower level of the hierarchy approaches steady-state. Multistable rivalry in this situation requires stronger mutual inhibition at the higher level of the model. However, if this is the case, changes in interocular grouping strength have the same effect on all the percepts. As a consequence Levelt’s propositions do not hold. We conclude that multistable rivalry is possible with inhibition only at the higher level of the visual hierarchy. However, mutual inhibition at the lower level is necessary for generalized Levelt’s propositions to hold.

Next we ask whether the mutual inhibition at the upper level is necessary for generalized Levelt’s propositions to be valid. Our model showed that it was not. The four propositions hold without mutual inhibition at the upper level (Fig. 6): The predominance of the (weaker) grouped percepts increases with *β* (Fig. 6A), and the average dominance duration of the (stronger) single-eye percepts decreases faster than that of the (weaker) grouped percepts increases (Fig. 6B). The average dominance duration of all percepts decreases as *α* = *β* increases (Fig. 6C).

Weak or mild mutual inhibition at the upper level does help improve the persistence of dominant percepts by increasing the difference between the activity levels of the dominant and suppressed percepts. Nonetheless, dominance switches still tend to be mainly determined by the activity at the lower level (See Fig. 7), as the dominance of a percept becomes increasingly clear as mutual inhibition is increased.

### 3.7. Impact of Adaptation at the Different Levels

Adaptation plays a central role in most models of rivalry, by decreasing the stability of the dominant percept, and thus driving transitions between percepts (Kang and Blake, 2010; Hollins and Hudnell, 1980; Roumani and Moutoussis, 2012; Blake and Overton, 1979; Blake et al., 1990; van Boxtel et al., 2008; Wade and Weert, 1986). We therefore asked at what level of the visual hierarchy this type of adaptation is needed to explain experimentally observed switching dynamics. As with mutual inhibition, we found that the generalized Levelt’s Propositions did hold when we removed adaptation (*κ* = 0) at the second Level of the population model (See Fig. 8). In addition, a change in the strength of adaptation had little effect on the average dominance of either grouped percepts or single-eye percepts. See Fig. 11A for example. However, when we removed adaptation at the lower level, the activity of lower level populations approached steady state since adaptation was necessary for switching to occur, and the generalized propositions did not hold any more.

### 3.8. Impact of Feedback

So far, we assumed an absence of feedback (*a*_*i*_ = 0 and *b*_*i*_ = 0) from the higher level of the visual hierarchy. However, numerous studies have found top–down feedback pathways from higher areas processing more complex features to lower areas processing basic geometric features (Angelucci et al., 2002; van Ee et al., 2006; Tong et al., 2006). Thus, we next asked whether generalized Levelt’s propositions still hold when we included feedback in our model as described in Eqs. (5a) -(5d). Our simulations showed that for weak feedback (*a*_*i*_ and *b*_*i*_ small), the dynamics of the hierarchical model described above did not change qualitatively (Compare Fig. 9 with feedback, to Fig. 4, with no feedback). However, the average dominance duration was larger when we included feedback, consistent with findings in the bistable case (Wilson, 2003).

### 3.9. The hierarchical model captures bistable binocular rivalry

As our hierarchical model is an extension of earlier models of binocular rivalry, we asked whether it also exhibits dynamics consistent with rivalry between two percepts. To answer this question we provided coherent “stimuli” to each pair of populations receiving input from the same eye, but conflicting stimuli to the two eyes. This would be equivalent to displaying a monochromatic square composed of vertical bars to one eye, and a monochromatic square composed of horizontal bars to the other eye.

Without feedback and including weak mutual inhibition and adaptation at the higher level, the dynamics of the system is mainly determined by that of the lower–level populations. Hence the only active populations at the higher level are therefore those corresponding to single–eye percepts. More precisely, without noise, and assuming *I*_1_ = *I*_2_, *I*_3_ = *I*_4_, the subsystem at the lower level has a flow-invariant subspace, *S* = {*E*_1_ = *E*_2_, *E*_3_ = *E*_4_, *H*_1_ = *H*_2_, *H*_3_ = *H*_4_}. Diekman et al. (2012) proved the subspace *S* is locally attracting at every point. When restricted to the subspace *S*, Eq. (1) reduces to a classical two population model (Laing and Chow, 2002; Wilson, 2003):

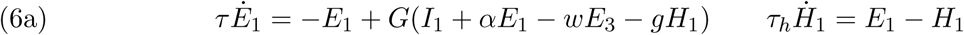

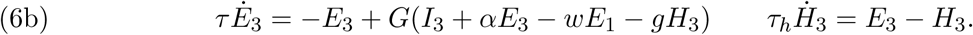

When population *E*_1_(= *E*_2_) dominates, it leads to the domination of percept 1 (*P*_1_). Similarly, when *E*_3_(= *E*_4_) dominates, then so does percept 2 (*P*_2_). Alternations in elevated activity between populations *E*_1_ and *E*_3_ therefore correspond to rivalry between percepts 1 and 2. Hence, Eq. (1) generalizes existing models of rivalry, and can capture features of binocular and multistable rivalry observed in experiments.

In addition, while the synchrony subspace *S* is associated with single-eye percepts (when *E*_1_ = *E*_2_ *> E*_3_ = *E*_4_, *P*_1_ dominates; when *E*_3_ = *E*_4_ *> E*_1_ = *E*_2_, *P*_2_ dominates), if *I*_1_ = *I*_4_, *I*_2_ = *I*_3_, then there is another synchrony subspace *W* = {*E*_1_ = *E*_4_, *E*_2_ = *E*_3_} (when *I*_1_ = *I*_4_, *I*_2_ = *I*_3_) associated to grouped percepts (when *E*_1_ = *E*_4_ *> E*_3_ = *E*_2_, *P*_3_ dominates; when *E*_3_ = *E*_2_ *> E*_1_ = *E*_4_, *P*_4_ dominates). The model thus also suggests that with sufficiently strong cues, the dynamics could be restricted to the invariant subset *W*, resulting in pure pattern rivalry.

## 4. Discussion

Multistable perceptual phenomena have long been used to probe the mechanisms underlying visual processing (Leopold and Logothetis, 1999). Among these, binocular rivalry is perhaps the most robust, and has been studied most frequently. However, we can obtain different insights by employing visual inputs that are integrated to produce interocularly grouped percepts (Kovacs et al., 1996; Suzuki and Grabowecky, 2002). These experiments are particularly informative when guided by Levelt’s Propositions, which were originally proposed to describe alternations between two rivaling percepts (Levelt, 1965; Brascamp et al., 2015).

We generalized Levelt’s Propositions to perceptual multistability involving interocular grouping. These extended propositions are consistent with experimental findings, and the dynamics of a hierarchical model of visual processing. Our neural population model thus points to potential mechanisms that underlie experimentally reported perceptual alternations in rivalry with interocular grouping (Jacot-Guillarmod et al., 2017).

Evidence suggests that rivalry exists across a hierarchy of visual cortical areas (Alias and Blake, 2004). Indeed, rivalry can occur between complex stimulus representations, requiring higher order processing than typically observed in early visual areas (Kovacs et al., 1996; Tong et al., 2006). Physiological and imaging experiments have also shown that binocular rivalry modulates neural activities in the primary visual cortex, as well as higher areas including V2 and V4, MT, and inferior temporal cortex (Leopold and Logothetis, 1996; Logothetis and Schall, 1989; Sheinberg and Logothetis, 1997; Tong and et al, 1998). However, the way in which activity at these different levels contributes to binocular rivalry remains unclear. Competition at the lower or higher levels, or a combination thereof can all explain different aspects of this phenomenon, depending on the experiment (Leopold and Logothetis, 1999; Pearson et al., 2007). Our model suggests that mutual inhibition at the early stages of the visual hierarchy is necessary for dynamics consistent with generalized Levelt’s Propositions.

Multistable rivalry has been studied previously using interocular grouping and fusion of coherently moving gratings. Moving plaid percepts arise when superimposing two drifting gratings moving at an angle to one another (Hupe and Rubin, 2004). In these cases subjects perceive either a grating or a moving plaid in alternation (three total percepts: moving to the left, moving the right and moving upward). Mutual inhibitory, adapting neuronal network models display dynamics consistent with data from such experiments, suggesting the mechanisms behind such rivalry may be similar to those driving conventional binocular rivalry (Huguet et al., 2014). This provides further evidence that the classical models of rivalry can serve as a foundation for models describing more complex settings.

### Comparisons with previous models of perceptual multistability

Our computational model is based on the assumption that perceptual multistability occurs via a winner-take-all process, with a single percept temporarily excluding all others (Wilson, 2003; Shpiro et al., 2007). Consequently, some neural process must allow the system to switch from the dominant percept to another after a few seconds (Laing and Chow, 2002). The simplest model of this process is a multistable system with slow adaptation and/or noise drives switches between multiple attractors (Moreno-Bote et al., 2007; Braun and Mattia, 2010). This framework is common in models of binocular rivalry (Laing and Chow, 2002; Shpiro et al., 2007), non-eye-based perceptual rivalry (Brascamp et al., 2009), and even perceptual multistability with more than two percepts (Diekman et al., 2013; Kilpatrick, 2013; Huguet et al., 2014). Each percept typically corresponds to a single neural population which mutually inhibits the other(s). Spike rate adaptation or short term plasticity then drive the slow switching between network attractors (Laing and Chow, 2002), and noise generates variation in the dominance times (Moreno-Bote et al., 2007).

Our computational model differs from previous ones in a few key ways. Excitatory connectivity at the first level facilitates both single-eye and grouped binocular percepts. Diekman et al. (2013) provided a preliminary account of interocular grouping, but ignored the effects of noise fluctuations on switching dynamics, and did not account for the known hierarchical structure of the visual system (Angelucci et al., 2002; Tong et al., 2006). In our model the strength of excitatory connectivity at the first level determines the input strength to populations at the higher level of the visual hierarchy, and ultimately each percept’s predominance. In this way, our model is similar to that in Wilson (2003), who used a two level model to capture the effects of monocular and binocular neurons. However, Wilson’s model focused on the case of two possible percepts, while our computational model accounts for all four possible percepts in an interocular grouping task.

A number of other hierarchical models have also been proposed: Dayan (1998) developed a top-down statistical generative model, which places the competition at the higher level. Freeman (2005) proposed a feedforward multistage model with all stages possessing the same structure. These models also focused on conventional bistable binocular rivalry, and did not address the mechanisms of multistable rivalry.

### Extensions to other computational models

We made several specific choices in our computational model. First, we described neural responses to input in each visual hemifield by a single variable. We could also have partitioned population activity based on orientation selectivity or receptive field location (Ferster and Miller, 2000). This would allow us to describe the effects of horizontal connections that facilitate the representation of collinear orientation segments in more detail (Bosking et al., 1997; Angelucci et al., 2002). Since there is evidence for chromatically-dependent collinear facilitation (Beaudot and Mullen, 2003), we could model the effects of image contrast and color saturation as separate contributions to interocular grouping. However, these extensions would complicate the model and make it more difficult to analyze. We therefore chose a reduced model with the effects of color saturation described by a single parameter, *β*.

### Neural mechanisms of perceptual multistability

Our observations support the prevailing theory that perceptual multistability is significantly percept-based and involves higher visual and object-recognition areas (Leopold and Logothetis, 1999). However, a number of issues remain unresolved. The question of whether and when binocular rivalry is eye-based or percept-based has not been fully answered (Blake, 2001). Activity predictive of a subject’s dominant percept has been recorded in lateral geniculate nucleus (LGN) (Haynes and Rees, 2005), primary visual cortex (V1) (Lee and Blake, 2002; Polonsky et al., 2000), and higher visual areas (e.g., V2, V4, MT, IT) (Logothetis and Schall, 1989; Leopold and Logothetis, 1996; Sheinberg and Logothetis, 1997). Thus, rivalry likely results from interactions between networks at several levels of the visual system (Freeman, 2005; Wilson, 2003). To understand how these activities collectively determine perception it is hence important to develop descriptive models that incorporate multiple levels of the visual processing hierarchy.

Collinear facilitation involves both recurrent connectivity in V1 as well as feedback connections from higher visual areas like V2 (Angelucci et al., 2002; Gilbert and Sigman, 2007), reenforcing the notion that perceptual rivalry engages a distributed neural architecture. However, a coherent theory that relates image features to dominance statistics during perceptual switching is lacking. It is unclear how neurons that are associated to each subpopulation may interact due to grouping factors such as collinearity and color.

### Conclusion

Our work supports the general notion that perceptual multistability is a distributed process that engages several layers of the visual system. Interocular grouping requires integration in higher visual areas (Leopold and Logothetis, 1996), but orientation processing and competition occurs earlier in the visual stream (Angelucci et al., 2002; Gilbert and Sigman, 2007). Overall, our model shows that the mechanisms that explain bistable perceptual rivalry can indeed be extended to multistable perceptual rivalry.

## 5. Acknowledgments

We thank Martin Golubitsky for helpful suggestions. Funding was provided by DHS-2014-ST-062-000057, NSF HRD-1800406 and NSF CNS-1831980 (YW); NSF DMS-1615737, NSF DMS-1853630 (ZPK); NIH-1R01MH115557 (KJ and ZPK); DBI-1707400 (KJ).

## Appendix A. Choice of parameter values

We had to set a number of parameters in our model to capture the perceptual alternations observed experimentally. To do so we first let *α* = *β*, and chose a set of parameter values so that the corresponding deterministic model had a periodic solution with *E*_1_(*t*) = *E*_2_(*t*) and *E*_3_(*t*) = *E*_4_(*t*). *i.e*. the periodic solution associated with the alternation of single-eye percepts. We then used XPPAUT to obtain the bifurcation diagram shown in Fig. 10, where the green curve in (A) is a branch of stable periodic solutions and the green curve in (B) is the corresponding periods of the periodic solutions in (A). We choose the values of input strength *I*_*i*_ all to be equal and in the interval (0.8, 1.25) so that the model displayed decreases in dominance duration with increasing input strength *I*.

Changing the values of *α* and *β* changes the bifurcation diagram. However, by continuity, as long as parameter values are not far from those we used to obtain the bifurcation diagram, the dynamics of the system remains similar. In many of our simulations, we fixed the input values *I* to 1.2, and other values at *α* = 0.3, *w* = 1, *g* = 0.5, *c*_*i*_ = 1, *v* = *γ* = 0.45, *κ* = 0.5. *τ* = 10*ms, τ*_*h*_ = *τ*_*a*_ = 1000*ms, δ* = 0.03. The parameter values of *w, g, v, γ* and *κ* roughly follow the values used in the literature (Seely and Chow, 2011; Wilson, 2003). We then numerically found the same qualitative results hold for *I ∈* [1, 1.25].

## Appendix B. Simulation procedure

To obtain the results shown in the figure, for each given parameter set we ran 100 realizations of the model for 300 seconds each and computed the dominance durations, predominance, and visit ratio for each percept. We pooled all dominance durations of one class of percepts (e.g., single-eye percepts or grouped percepts) and computed its average and standard deviation across occurrences and realizations.

## Appendix C. Simulation results with feedback from higher to lower level

Our hierarchical model with sufficiently weak feedback from the higher level to the lower level can also capture the three main observations reported by Jacot-Guillarmod et al. (2017) with the minor difference that the average dominance duration increases (Fig. 9). Increasing the adaptation rate *κ* in the top level had little or no effect on the dominance duration of percepts (Fig. 11A shows single-eye percepts, but results for grouped percepts were similar) over a large interval (0, 0.8). The main effect of top down excitatory feedback from a percept we observed was to increase that percept’s dominance duration (Fig. 11B).

## References

Alias, D., Blake, R., 2004. Binocular Rivalry. MIT Press, Cambridge, MA.

Andrew, T. J., Lotto, R. B., 2004. Fusion and rivalry are dependent on the perceptual meaning of visual stimuli. Curr. Biol. 14, 418–423.

Angelucci, A., Levitt, J. B., Walton, E. J., Hupe, J.-M., Bullier, J., Lund, J. S., 2002. Circuits for local and global signal integration in primary visual cortex. The Journal of Neuroscience 22 (19), 8633–8646.

Arrington, K. F., 1993. Neural network models for color brightness percception and binocular rivalry. Ph.D. thesis, Boston University.

Bartels, A., Logothetis, N. K., 2010. Binocular rivalry: a time-depedence of eye and stimulus contributions. Journal of vision 10, 1–14.

Beaudot, W. H., Mullen, K. T., 2003. How long range is contour integration in human color vision? Visual Neuroscience 20 (01), 51–64.

Benda, J., Herz, A. V., 2003. A universal model for spike-frequency adaptation. Neural computation 15 (11), 2523–2564.

Blake, R., 1989. A neural theory of binocular rivalry. Psychol. Rev. 96, 145–167.

Blake, R., 2001. A primer on binocular rivalry, including current controversies. Brain and Mind 2, 5–38.

Blake, R., Logothetis, N. K., 2002. Visual competition. Nat. Rev. Neurosci 3, 13–21.

Blake, R., Overton, R., 2019/09/27 1979. The site of binocular rivalry suppression. Perception 8 (2), 143–152. URL https://doi.org/10.1068/p080143

Blake, R., Westendorf, D., Fox, R., Nov 1990. Temporal perturbations of binocular rivalry. Perception & Psychophysics 48 (6), 593–602. URL https://doi.org/10.3758/BF03211605

Bosking, W., Zhang, Y., Schofield, B., Fitzpatrick, D., 1997. Orientation selectivity and the arrangement of horizontal connections in tree shrew striate cortex. Journal of neuroscience 17, 2112–2127.

Brascamp, J., Klink, P., Levelt, W. J., 2015. The ‘laws’ of binocular rivalry: 50 years of levelt’s propositions. Vision research 109, 20–37.

Brascamp, J., Pearson, J., Blake, R., Van Den Berg, A., 2009. Intermittent ambiguous stimuli: Implicit memory causes periodic perceptual alternations. Journal of Vision 9 (3), 3.

Brascamp, J. W., Van, R. E., Noest, A. J., Jacobs, R. H., van den Berg, A. V., 2006. The time course of binocular rivalry reveals a fundamental role of noise. J. Vis. 6, 1244 – 1256.

Braun, J., Mattia, M., 2010. Attractors and noise: twin drivers of decisions and multistability. Neuroimage 52 (3), 740–751.

Brincat, S. L., Connor, C. E., 2006. Dynamic shape synthesis in posterior inferotemporal cortex. Neuron 49 (1), 17–24.

Carlson, T. A., He, S., 2004. Competing global representations fail to initiate binocular rivalry. Neuron 43, 970–914.

Curtu, R., Shpiro, A., Rubin, N., Rinzel, J., 2008. Mechanisms for frequency control in neuronal competition models. SIAM J. Appl. Dyn. Sys. 7, 609–649.

Dayan, P., 1998. A hierarchical model of binocular rivalry. Neural Comput. 10, 1119–1135.

Diekman, C. O., Golubitsky, M., McMillen, T., 2012. Reduction and dynamics of a generalized rivalry network with two learned patterns. SIAM J. Appl. Dyn. Sys. 11, 1270–1309.

Diekman, C. O., Golubitsky, M., Wang, Y., 2013. Derived patterns in binocular rivalry networks. Journal of Mathematical Neuroscience 3 (6).

Faisal, A. A., Selen, L. P. J., Wolpert, D. M., Apr 2008. Noise in the nervous system. Nat Rev Neurosci 9 (4), 292–303.

Ferster, D., Miller, K. D., 2000. Neural mechanisms of orientation selectivity in the visual cortex. Annual review of neuroscience 23 (1), 441–471.

Freeman, A. W., 2005. Multistage model for binocular rivalry. J. Neurophysiol 94, 4412–4420.

Gilbert, C. D., Sigman, M., 2007. Brain states: top-down influences in sensory processing. Neuron 54 (5), 677–696.

Golubitsky, M., Zhao, Y., Wang, Y., Lu, Z.-L., 2019. The symmetry of generalized rivalry network models determines patterns of interocular grouping in four-location binocular rivalry. Journal of Neurophysiology.

Häusser, M., Roth, A., 1997. Estimating the time course of the excitatory synaptic conductance in neocortical pyramidal cells using a novel voltage jump method. The Journal of neuroscience 17 (20), 7606–7625.

Haynes, J. D., Rees, G., 2005. Predicting the stream of consciousness from activity in human visual cortex. Curr. Biol. 15 (14), 1301–7.

Hollins, M., Hudnell, K., 09 1980. Adaptation of the binocular rivalry mechanism. Investigative Ophthalmology & Visual Science 19 (9), 1117–1120.

Huguet, G., Rinzel, J., Hupé, J. M., 2014. Noise and adaptation in multistable perception: Noise drives when to switch, adaptation determines percept choice. J Vis 14 (3).

Hupe, J.-M., Rubin, N., 2004. The oblique plaid effect. Vision Research 44 (5), 489–500.

Jacot-Guillarmod, A., Wang, Y., Pedroza, C., Ogmen, H., Kilpatrick, Z., Josić, K., 2017. Extending levelt’s propositions to perceptual multistability involving interocular grouping. Submitted.

Kalarickal, G. J., Marshall, J. A., 2000. Neural model of temporal and stochastic properties of binocular rivalry. Neurocomputing 32, 843–853.

Kang, M.-S., Blake, R., Jan 2010. What causes alternations in dominance during binocular rivalry? Atten Percept Psychophys 72 (1), 179–86.

Kilpatrick, Z. P., 2013. Short term synaptic depression improves information transfer in perceptual multistability. Front Comput Neurosci 7 (85).

Klink, P. C., Brascamp, J. W., Blake, R., Wezel, R. J. A. V., 1464-1469 2010. Experience-driven plasticity in binocular vision. Curr. Biol. 20.

Klink, P. C., van Ee, R., Nijs, M. M., Brouwer, G. J., Noest, A. J., van Wezel, R. J. A., 2008. Early interacions between neuronal adaptation and voluntary control determine perceptual choices in bistable vision. Journal of Vision 8 (16), 1–18.

Kohler, W., 2015. The task of Gestalt psychology. Princeton University Press.

Kovacs, I., Papathomas, T. V., Yang, M., Feher, A., 1996. When the brain changes its mind: Interocular grouping during binocular rivalry. PNAS 93, 15508–15511.

Lago-Fernandez, L. F., Deco, G., 2002. A model of binocular rivalry based on competition in it. Neurocomputing 44-46, 503–507.

Laing, C., Chow, C. C., 2002. A spiking neuron model for binocular rivalry. J. Comput. Neurosci. 12, 39–53.

Lamme, V. A., Roelfsema, P. R., 2000. The distinct modes of vision offered by feedforward and recurrent processing. Trends in neurosciences 23 (11), 571–579.

Lankheet, M. J. M., 2006. Unraveling adaptation and mutual inhibition in perceptual rivalry. Journal of Vision 6, 304–310.

Lee, S.-H., Blake, R., 2002. V1 activity is reduced during binocular rivalry. Journal of Vision 2 (9), 4.

Lehky, S. R., 1988. An astable multivibrator model of binocular rivalry. Perception 17, 215–228.

Leopold, D., Logothetis, N. K., 1999. Multistable phenomena: changing views in perception. Trends in Cognitive Sciences 3, 254–264.

Leopold, D. A., Logothetis, N. K., 1996. Activity changes in early visual cortex refect monkeys’ percepts during binocular rivalry. Nature 379, 549–553.

Levelt, W. J. M., 1965. On binocular rivalry. Ph.D. thesis, Institute for Perception RVO-TNO Soeterberg (Netherlands).

Logothetis, N. K., Schall, J. D., 1989. Neuronal correlates of subjective visual perception. Science 245 (4919), 761–763.

Lumer, E. D., 1998. A neural model of binocular intergration and rivalry based on the coordination of action-potential timing in primary visual cortex. Cereb. Cortex 8, 553–561.

Matsuoka, K., 1984. The dynamic model of binocular rivalry. Biol Cybern 49, 201–208.

Moreno-Bote, R., Rinzel, J., Rubin, N., 2007. Noise-induced alternations in an attractor network model of perceptual bistability. Journal of Neurophysiology 98, 1125–1139.

Moreno-Bote, R., Shapiro, A., Rinzel, J., Rubin, N., 2010. Alternation rate in perceptual bistability is maximal at and symmetric around equi-dominance. Journal of vision 10 (11), 1–18.

Noest, A. J., van Ee, R., Nijs, M. M., van Wezel, R. J. A., 2007. Percept-choice sequences driven by interrupted ambiguous stimuli: a low-levl neural model. Journal of Vision 7 (8), 1–14.

Pearson, J., Tadin, D., Blake, R., May 2007. The effects of transcranial maganetic stimulation on visual rivalry. Journal of Vision 7 (7), 1–11.

Polonsky, A., Blake, R., Braun, J., Heeger, D., 2000. Neuronal activity in human primary visual cortex correlates with perception during binocular rivalry. Nat. Neurosci 3, 1153–1159.

Renart, A., De La Rocha, J., Bartho, P., Hollender, L., Parga, N., Reyes, A., Harris, K. D., 2010. The asynchronous state in cortical circuits. Science 327 (5965), 587–590.

Riesenhuber, M., Poggio, T., Nov. 1999. Hierarchical models of object recognition in cortex. Nature Neuroscience 2 (11), 1019–1025.

Ringach, D. L., Hawken, M. J., Shapley, R., et al., 1997. Dynamics of orientation tuning in macaque primary visual cortex. Nature 387 (6630), 281–284.

Roelfsema, P. R., 2006. Cortical algorithms for perceptual grouping. Annu. Rev. Neurosci. 29, 203–227.

Roumani, D., Moutoussis, K., 03 2012. Binocular rivalry alternations and their relation to visual adaptation. Frontiers in human neuroscience 6, 35–35. URL https://www.ncbi.nlm.nih.gov/pubmed/22403533

Said, C. P., Heeger, D. J., 2013. A model of binocular rivalry and cross-orientation suppression. Plos Computational Biology 9, e1002991.

Salinas, E., Abbott, L., 1996. A model of multiplicative neural responses in parietal cortex. Proceedings of the national academy of sciences 93 (21), 11956–11961.

Seely, J., Chow, C. C., 2011. The role of mutual inhibition in binocular rivalry. J Neurophysiol 106, 2136–2150.

Sheinberg, D. L., Logothetis, N. K., 1997. The role of temporal cortical areas in perceptual organization. Proceedings of the National Academy of Sciences 94 (7), 3408–3413.

Shpiro, A., Curtu, R., Rinzel, J., Rubin, N., 2007. Dynamical characteristics common to neuronal competition models. Journal of neurophysiology 97 (1), 462–473.

Sincich, L. C., Horton, J. C., 2005. The circuitry of v1 and v2: integration of color, form, and motion. Annu. Rev. Neurosci. 28, 303–326.

Sterzer, P., Kleinschmit, A., Rees, G., 2009. The neural bases of multistable perception. Trends in Cognitive Sciences 13 (7).

Stettler, D. D., Das, A., Bennett, J., Gilbert, C. D., 2002. Lateral connectivity and contextual interactions in macaque primary visual cortex. Neuron 36 (4), 739–750.

Stollenwerk, L., Bode, M., 2003. Lateral neural model of binocular rivalry. ftrNeural Comput. 15, 2863–2882.

Suzuki, S., Grabowecky, M., 2002. Evidence for perceptual “trapping” and adaptation in multistable binocular rivalry. Neuron 36, 143–157.

Tong, F., 2001. Competing theories of binocular rivalry. Brain and Mind 2, 55083.

Tong, F., et al, 1998. Binocular rivalry and visual awareness in human extrastriate cortex. Neuron 21 (753–759).

Tong, F., Meng, M., Blake, R., 2006. Neural bases of binocular rivalry. Trends in Cognitive Sciences 10 (11).

van Boxtel, J. J. A., Alais, D., van Ee, R., 05 2008. Retinotopic and non-retinotopic stimulus encoding in binocular rivalry and the involvement of feedback. Journal of Vision 8 (5), 17-17. URL https://doi.org/10.1167/8.5.17

van Ee, R., Noest, A. J., Brascamp, J. W., van den Berg, A. V., 2006. Attentional control over either of the two competing percepts of ambiguous stimuli revealed by a two-parameter analysis: Means do notmake the difference. Vision Research 46, 3129–3141.

Wade, N. J., Weert, C. M. M. D., 1986. Aftereffects in binocular rivalry. Perception 15 (4), 419–434, pMID: 3822728. URL https://doi.org/10.1068/p150419

Wagemans, J., Elder, J. H., Kubovy, M., Palmer, S. E., Peterson, M. A., Singh, M., von der Heydt, R., 2012. A century of gestalt psychology in visual perception: I. perceptual grouping and figure–ground organization. Psychological bulletin 138 (6), 1172.

Wheatstone, C., 1838. Contributions to the physilogy of vision. part i. on some remakable, and hitherto unobserved, phenomena of bioncular vision. London and Edinberugh Philosophical Magazing and Journal of Science 3, 241–267.

Wilson, H. R., 2003. Computational evidence for a rivalry hierarchy in vision. PNAS 100, 14499–14503.

Wilson, H. R., 2007. Minimal physiological conditions for binocular rivalry and rivalry memory. Vision Res 47, 2741–2750.

Wilson, H. R., 2009. Requirements for conscious visual processing. Cambridge University Press, Ch. In Cortical Mechanisms of Vision, pp. 399–417.

